# Triazole 187 is a biased KOR agonist that suppresses itch without sedation and induces anxiolytic-like behaviors in mice

**DOI:** 10.1101/2025.02.17.638680

**Authors:** Allison Volf, Tarsis F. Brust, Robin R. Kobylski, Kerri M. Czekner, Edward L. Stahl, Michael D. Cameron, Ashley E. Trojniak, Jeffrey Aubé, Laura M. Bohn

## Abstract

Kappa opioid receptor agonists are clinically used to treat pruritis and have therapeutic potential for the treatment of pain and neuropsychiatric disorders. We have previously shown that triazole 1.1 is a G protein signaling-biased KOR agonist, that can suppress itch without producing signs of sedation in mice. This profile was recapitulated in rats and non-human primates however, triazole 1.1 had limited potency as an antipruritic. Here we describe a more potent, G protein signaling-biased agonist, triazole 187. Triazole 187 is a potent antipruritic agent and does not decrease spontaneous locomotor activity; interestingly, it produces anxiolytic-like behaviors in mice, an effect not observed for triazole 1.1. In addition to curbing sedation, triazole 187 produces only mild diuresis, resulting in 30% of urine output induced by U50,488H at dose that is 188-fold the antipruritic potency dose. Compounds like triazole 187 may present a means to treat anxiety accompanied by persistent chronic itch while avoiding sedation and diuresis accompanied by typical KOR agonists.

Graphic Abstract

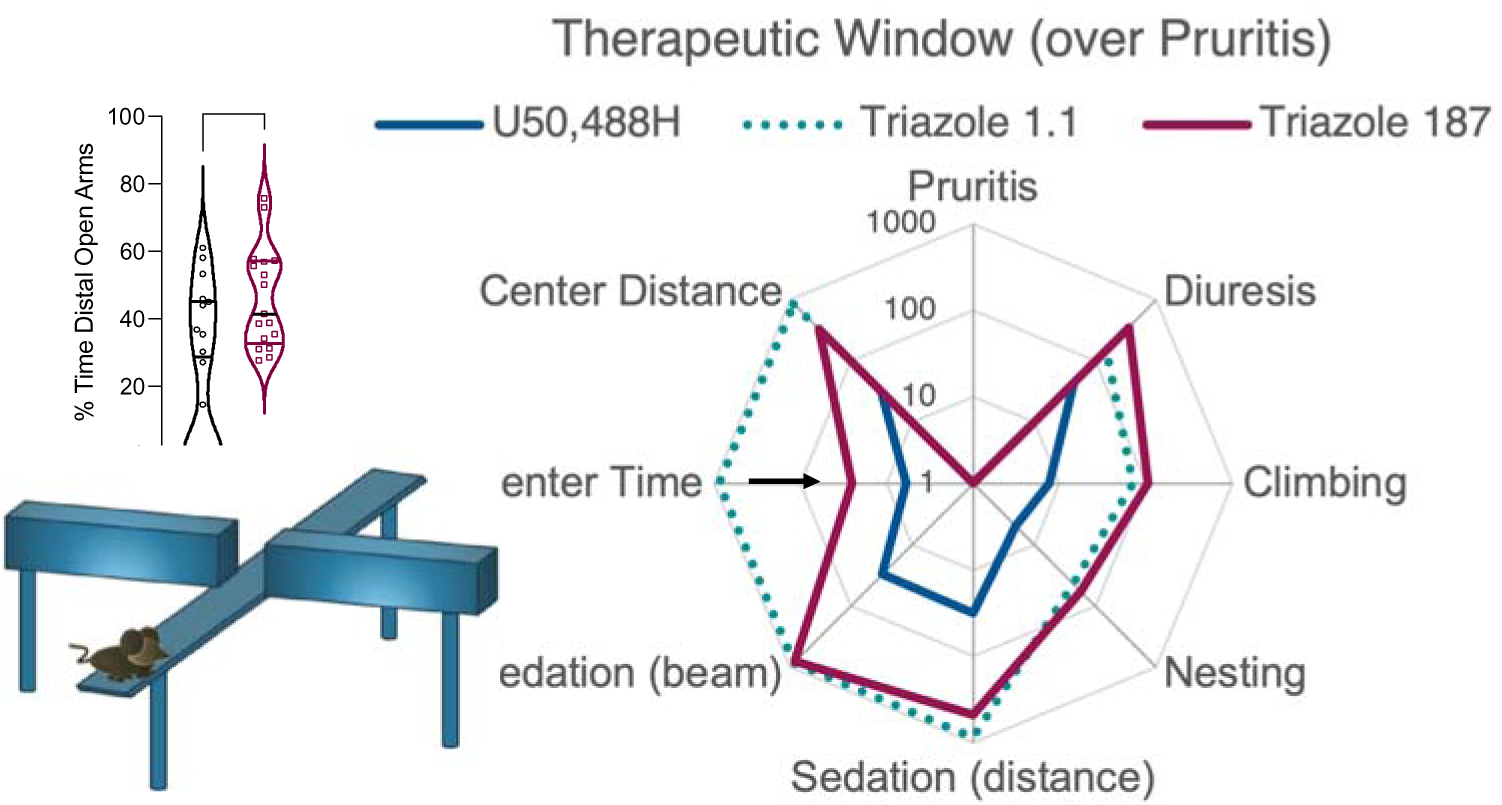

## INTRODUCTION

The kappa opioid receptor (KOR) is a seven-transmembrane spanning G protein-coupled receptor (GPCR)^1^ distributed throughout the central and peripheral nervous system (CNS and PNS)^2^. KOR agonists are clinically used for the treatment of non-histamine-induced itch, or pruritis^3–7^, while also being pursued as a target in therapies such as pain^8, 9^, anxiety^10–12^, opioid abuse disorder^13–16^ and as potential non-serotonergic psychedelic therapy^17, 18^. Unfortunately, activation of KOR in the central nervous system (CNS) can lead to undesired side effects, such as diuresis, sedation, and dysphoria^19–24^ which has limited the clinical utility of KOR agonists.

To avoid CNS-mediated side effects, drug development has focused on KOR agonists that are restricted to the periphery as a treatment for pruritus. These efforts have led to the FDA-approval of difelikefalin to treat pruritus associated with chronic kidney disease in adults undergoing hemodialysis^3^. While dysphoria and hallucinations have not been reported by patients treated with recommended doses of difelikefalin; drowsiness, diarrhea, and dizziness have been noted despite the “peripheral restriction” of the drug^25^. The KOR agonist, nalfurafine, is the only brain penetrant KOR agonist clinically available for the treatment of pruritus and, at the time of writing, is only approved in Japan^4, 5, 26^. At clinically relevant doses in humans, nalfurafine produces anti-pruritic effects without causing sedation or dysphoria^5, 27^, however side effects such as drowsiness and constipation have been observed^26^. These side effects may be attributed to nonselectivity as nalfurafine is also an agonist at other receptors, such as the mu opioid receptor^5^. Interestingly, nalfurafine has revealed atypical KOR agonist pharmacology and has been described as a G protein-signaling biased agonist^5^.

Biased agonism has been explored as a means to optimize therapeutic properties while avoiding certain side effects^28, 29^. We previously introduced triazole 1.1^30^ and have shown that this fully efficacious agonist preferentially promotes KOR-mediated ^35^S-GTPγS binding over βarrestin2 recruitment, adenylyl cyclase inhibition, and ERK1/2 activation relative to conventional KOR agonists U69,593 or U50,488H^22, 31, 32^. Triazole 1.1 is brain penetrant and binds to KOR with high selectivity^22, 30^. In mice, triazole 1.1 is antipruritic and antinociceptive, but unlike typical KOR agonists, such as U50,488H, triazole 1.1 does not decrease spontaneous locomotor activity^22^. Such dose-dependent decreases in ambulatory behavior in mice has been shown to correlate with sedative-like drug effects in higher species^33–35^ suggesting that triazole 1.1 may not be sedating. When tested in non-human primates, triazole 1.1 produces antinociception without sedation; moreover, when given with oxycodone, triazole 1.1 suppresses opioid-induced itch^16, 36^. However, higher doses of triazole 1.1 are necessary to achieve the same antipruritic effect as U50,488H, indicating its potency could be improved upon^22, 36^.

In this study, we evaluated triazole 187, a member of a new series of triazole agonists^37^. With the replacement of a thioether and a furan group by a carbon and a thiophenyl respectively, we demonstrate in cell signaling assays that triazole 187 shows improved potency in multiple signaling cascades, while retaining biased agonism for promoting ^35^S-GTPγS binding over βarrestin2 recruitment, adenylyl cyclase inhibition, and ERK1/2 activation. Like triazole 1.1, triazole 187 enters the brain when administered systemically and binds to KOR with high selectivity. Herein we present the potency and efficacy of triazole 187, U50,488H and triazole 1.1 in several mouse models of KOR activity. Overall triazole 187 has improved antipruritic potency compared to triazole 1.1 while avoiding sedation and limiting the extent of diuresis. Triazole 187 produces anxiolytic-like behaviors that are not confounded by sedative properties typically seen in KOR agonists. As pruritis can be accompanied by negative anticipations and anxiety, KOR agonists that can enter the brain and positively affect mood might provide a means to not only treat itch, but also treat the anxiety that accompanies intractable itch.

## MATERIALS AND METHODS

### Chemicals and drugs

In this study, the following compounds were purchased as indicated: U69,593 from Sigma-Aldrich (St. Louis, MO); U50,488H from Tocris (Ellisville, MO); Tween 80 and dimethyl sulfoxide (DMSO) (certified grade reference material pharmaceutical secondary standard, >=99.7% purity) from Fisher Scientific (Pittsburg, PA); and sterile 0.9% saline from VWR (Radnor, PA). Triazole 1.1 ^30^ and triazole 187 ^37^ were synthesized as previously described and were at >99% purity for all studies as determined by high performance liquid chromatography (HPLC). For all in vitro assays, compounds were prepared in DMSO at concentrations spanning from 32 nM to 10 mM, for dilutions with the final DMSO concentration of less than 1% in any assay.

### Cell lines and cell culture

Chinese hamster ovary (CHO) cells expressing human kappa opioid receptor (hKOR) complementary DNA (cDNA) including three haemagglutinin tags on the N-terminus (3×HA-hKOR, cDNA.org) were generated and used as previously described; the DiscoveRx PathHunter^TM^ U2OS hKOR (U2OS-hKOR-βarrestin2-EFC) were purchased from DiscoveRx Corp. (now MilliporeSigma, Burlington, MA) and are used as previously described^22, 30–32, 37, 38^.

### ^35^S-GTPγS binding assay

Cell membranes are prepared from CHO-hKOR cells and assessed for [^35^S]-GTPγS binding following treatment with KOR agonists according to previously published protocols^30, 32^. Briefly, 15 μg membranes were incubated in assay buffer (50 mM Tris pH 7.4, 100 mM NaCl, 5 mM MgCl_2_, 1 mM EDTA, 3 μM GDP) containing 0.1 nM [^35^S]GTPγS from Revvity, (Waltham, MA) (with specific activity of 1250 Ci/mmol) with agonist at a total reaction volume of 200 μl for 1 hour at 25 °C. Termination of the reaction is achieved by rapid filtration and washing with ice cold water through a GF/B glass microfiber filter (Whatman^®^ glass microfiber filters, Grade GF/B from Sigma Aldrich) using a 96-well plate harvester from Brandel Inc. (Gaithersburg, MD). Filters were punched into shallow 96-well plates compatible with the TopCount NXT Microplate Scintillation and Luminescence Counter from PerkinElmer Life Sciences (Waltham, MA) and air-dried filters are counted following addition of scintillation fluid (Microscint^TM^-20, from Revvity). Two technical replicates were included in each individual experiment (n≥ 3).

### Forskolin-stimulated cAMP accumulation assay

The CISBIO homogeneous time-resolved florescence (HTRF) cAMP kit (CISBIO HTRF cAMP Gs HiRange from Revvity) was used to assess cAMP levels as previously described in detail^32^. HTRF ratios (fluorescence at 665 nm/ fluorescence at 620 nm) were determined using a BioTek Synergy Neo2 Hybrid Multimode reader (Agilent, Santa Clara, CA) as a measure of inhibition of forskolin (20 µM) cAMP accumulation. Three technical replicates were included in each individual experiment (n≥ 3).

### βArrestin2 recruitment assay

βArrestin2 recruitment is determined using the DiscoveRx PathHunter enzyme complementation assay from MilliporeSigma in PathHunter U2OS OPRK1 βarrestin-2 cells according to manufacturer’s protocol and as previously described^30, 32, 38^. Luminescence values are determined by using a BioTek Synergy Neo2 Hybrid Multimode reader. All compounds were run in triplicate per assay and normalized to vehicle-treated cells; three technical replicates were included in each individual experiment (n≥ 3).

### In-cell western ERK1/2 phosphorylation

In-cell western ERK1/2 phosphorylation assays are performed according to previously published protocol^30, 38, 39^. Permeabilized cells are incubated with primary antibodies for phosphorylated ERK1/2 (1:300, Cell Signaling (Bevelry, MA) #4370) and total ERK1/2 (1:400 mouse, Cell Signaling #4696) at 4°C overnight in blocking buffer. After washing, secondary antibodies (Li-Cor; anti-rabbit IRDye800CW, 1:500; anti-mouse IRDye680LT, 1:1500 from LICORbio^TM^ (Lincoln, NE)) are then added in 1:1 Li-Cor blocking buffer in PBS, containing 0.025% Tween20 for 1 hour at 25 °C. Florescence is measured using an Odyssey M Imaging System from LICORbio^TM^ at 700 and 800 nm. The fold ERK1/2 phosphorylation over vehicle levels was determined by first normalizing phosphorylated ERK1/2 levels to total ERK1/2 levels within each well and then normalizing to the average of the vehicle treated cells within each experiment; three technical replicates were included in each individual experiment (n≥ 3).

### Radioligand binding assays

Membrane preparations are made after CHO-hKOR cell homogenization and subjected to radioligand binding assays according to a modified procedure to previously published protocols^30, 40^. Membranes (20 μg protein/well) are incubated for 2 hours at 25°C in the presence of 2 nM ^3^H-Naloxone (Revvity, USA, Concentration=1000 μCi/mL, SA=79.9 Ci/mmol) and test compounds with a final DMSO concentration of 1%. Radioligand binding assays are terminated by filtration through GF/B glass microfiber filters using a 96-well Brandel Cell Harvester followed by several washes with cold 10mM Tris (pH 7.4). Air-dried filters are counted on the Topcount plate reader following addition of scintillation fluid with 2.7-fold efficiency correction for converting counts per minute (CPM) to disintegrations per minute (DPM). Nonspecific binding was determined for each radioligand by competition with 1 μΜ naloxone. The affinity of ^3^H-Naloxone was determined by homologous competition: K_D_ = 3.6 nM. Competitive radioligand binding studies are analyzed using the heterologous competition nonlinear competition in GraphPad Prism to determine pK*_i_*; two technical replicates were included in each individual experiment (n≥ 3).

### Animals

Male C57BL/6J mice were purchased from The Jackson Laboratory. KOR-KO mice were purchased from The Jackson Laboratory and propagated using homozygous breeding as previously described^41^. Adult mice are used between 10 and 24 weeks of age and are only used once per assay. Mice are kept on a 12-hour light/dark cycle in a temperature-controlled room and are group-housed (three to five mice per cage). All behavioral tests were performed during the light cycle between 8am-6pm. For drug treatments mice are injected with compound or vehicle at 10 μl/g body weight. Injections are either subcutaneous (s.c.) or intraperitoneal (i.p.), indicated in figures and figure legends. Experimenters conducting injections and performing measurements are blinded to the treatment groups. The vehicle for animal studies is DMSO, Tween 80, and 0.9% sterile saline (1:1:8). To facilitate solubility, the compounds are prepared by dissolving drug powder in DMSO, adding warm (50°C) Tween80, then 0.9% saline and immediately vortexing; solutions were made and used immediately before each study. All experiments conducted with mice are approved by the Institutional Animal Care and Use Committee at The Herbert Wertheim UF Scripps Institute for Biomedical Innovation & Technology (Jupiter, FL) and adhere to the National Institutes of Health Animal Care guidelines.

### Pharmacokinetics

To determine drug levels in plasma and brain; male C57BL6/J mice were injected (i.p) with KOR agonists; plasma was taken at the indicated time points in figures. For brain levels, following cervical dislocation, brains were taken at 30, 60, 120, or 240 minutes and frozen in liquid nitrogen. Samples were subjected to Liquid chromatography (Shimadzu)–tandem mass spectrometry from AB Sciex (Framingham, MA) operated using multiple reaction monitoring with a mass transition of 429.3→97.1 Da for triazole 187, and 431.3→173.2 Da for triazole 1.1. Pharmacokinetic parameters were calculated using a noncompartmental model (Phoenix WinNonlin, Pharsight Inc.).

### Pruritus

The mouse pruritus assay was conducted as previously described^22, 42^. Mice are briefly acclimated to clear acrylic testing boxes (10 × 10 cm^2^) for 1 hour then injected with vehicle, U50,488H, triazole 1.1 or triazole 187 (injection indicated in figures) and returned to the acrylic boxes and video recording is started. Ten minutes after pretreatment, mice receive an injection of 40 mg/kg chloroquine phosphate (CP) from Sigma-Aldrich. CP is freshly prepared in a solution consisting of DMSO, Tween80 and 0.9% saline (1:1:8); first dissolved in 0.9% sterile saline and brought up to volume with DMSO and Tween80 to; pH 6.0 and injected s.c. in the skin at the base of the neck at a volume of 5 μl/g body weight. Mice are immediately placed back into the acrylic boxes and the number of scratching bouts is counted for one hour by an investigator blinded to the treatment groups. During the experiments, mice are videotaped and another investigator repeated scoring to validate method. One “scratching bout” is defined as one or more rapid movements of the hind paw toward the injection site before placing the paw back on the floor^43^. To conserve animals, the values obtained for vehicle (s.c.) and 1 mg/kg of U50,488H (s.c.) and 1 mg/kg triazole 1.1 (s.c.) are taken from the 2016 Brust et al. study^22^ as this was the lowest dose tested and used to establish the maximally efficacious dose.

### Diuresis

Male C57Bl/6J mice are habituated in the experimental room for 1 hour prior to experiment with access to water, after which they are handled and weighed. The investigators are blinded to treatments and doses, and the mice are injected s.c. with KOR agonist or vehicle and placed in 1/3 of a new (home) cage (18 cm x 12 cm x 12 cm) (separated with a divider) with the bedding replaced by hydrophobic sand (LabSand, Coastline Global, Inc., West Chester, PA). Subcutaneous administration was used in this assay for all compounds to prevent unintended pressure to the bladder. Mice are monitored for the duration of the test and urine is collected from the sand as soon as the mouse urinates using a 200 μL pipettor and dispensed into Eppendorf ( 1.5 mL) tubes which were closed to prevent evaporation. Final volumes were quantified using a 200 μL or 1 mL pipettor at the end of the test.

### Climbing behavior

During the diuresis assay, investigators observed mice climbing to the top of the home cage; these attempts were recorded. An attempt is defined as every time a mouse tried to climb out of its cage during the 3 hour study and experimenter had to direct the mouse back into its cage. If a mouse reached for the top of the cage wall but did not bring its body to the top it is not considered an attempt.

### Locomotor activity

The Versamax Animal Activity Monitoring System (20 cm × 20 cm x 30.5 cm) from Accuscan Instruments (Columbus, OH) was used to assess open-field locomotor activity as previously described^22, 42^. The system consists of a photocell-equipped automated open field chamber contained inside sound-mitigating boxes to record locomotor activity. The Versadat software from Accuscan Instruments records the overall activity (distance traveled, beam breaks, center time, and center distance) in 5-minute intervals. Without habituation, mice are injected with vehicle, U50,488H, triazole 1.1 or triazole 187 (injection indicated in figures) and placed immediately into the locomotor motor activity box after injection to record spontaneous locomotor activity over 60 minutes. To conserve animals, the data for vehicle, triazole 1.1 and U50,488H from the locomotor activity studies from the 2016 Brust et al., study^22^ were re-analyzed for total distance traveled, center time spent, and center distance. Only horizontal beam breaks were presented in the prior publication. To capture the potency of U50,488H, we added 10 mice at 1 mg/kg (not tested previously), 2 mice at 3 mg/kg, 4 mice at 5 mg/kg, 2 mice at 15 mg/kg, 4 mice at 30 mg/kg and 4 vehicle treated mice (s.c.) to the data analyzed from the original study.

### Mouse nesting

The mouse nesting assay was conducted similarly to previously described^44^. Briefly, mice are singly housed in a new (home) cage for three days for habituation before the start of the nesting study. Following habituation, mice are subject to 2 days of nesting sessions (one per day) to acclimate them to the experimental conditions and the procedure room. A nesting session entails briefly removing the mouse from the home cage and placing nestlets into 6 zones of the home cage. The mouse is then returned to the cage and an investigator measures the number of zones cleared over 1 hour. Clearing of a zone occurs when the nestlet is moved (usually to make one pile in a corner)^44^. On day 3 mice are given saline, placed in their home cage for 10 minutes, briefly removed to allow for placing of the nestlets in 6 zones, then returned to the home cage for nesting. Mice that failed to clear more than 3 nestlets after the saline injection were not used for further analysis. On day 4, investigators are blinded to treatments where mice are injected with vehicle, U50,488H, triazole 1.1, or triazole 187; photographs were collected and another blinded investigator also contributed to validating scoring.

### Elevated plus maze

Mice are placed into a structure shaped like a “plus sign”, in which two of the arms are exposed and two are enclosed with walls (black maze with white flooring; Med Associates, St. Albans, VT) as previously described^45^. Mice are injected (i.p) with indicated doses of KOR agonists or vehicle and returned to their home cages for 30 minutes before placing in the center of the plus maze and allowed to explore the maze for 5 minutes. Time spent in proximal and distal areas of each arm is electronically recorded for five minutes using EthoVision XT video recording from Noldus Information Technology Inc. (Leesburg, VA). Mice are excluded if they fell off the maze or if total activity is more than two standard deviations less than mean vehicle-treated activity. The elevated plus maze was performed by the Animal Behavior Core at The Herbert Wertheim UF Scripps Institute for Biomedical Innovation & Technology and the investigator was blinded to treatment. Normalization of time spent is based on the 5 minute test duration.

### Statistical and data analyses (in vitro)

All statistical comparisons are made using GraphPad Prism 10 software (GraphPad Software Inc., San Diego, CA) and is expressed as means ± SEM, unless indicated otherwise. For in vitro cell-based functional assays, agonist stimulation is determined and presented by normalizing all values to the top of the maximum response (E_max_) produced by U69,593 and using the baseline within the assay to determine the bottom. The values of half maximal effective concentration from individual experiment and (pEC_50_) and E_max_ values are derived from a three-parameter, nonlinear regression analysis on the experimental replicates and are presented as the mean with 95% CI in Table 1. A comparison of potency (logEC50) was performed within Prism using the extra sum-of-squares F test (p=0.05) to compare the curves derived from of at least three independent experiments per agonist. Biased agonism calculations are described below. For radioligand binding assays, pK*_i_*values are compared between U69,593 and each triazole using an extra sum-of-squares F test using GraphPad Prism to compare the global fit values derived from the curve of at least three independent experiments for each agonist. Individual experiments were performed in duplicate and normalized to vehicle-treated cells; data are presented as means of 3 or more individual experiments (mean with S.E.M. data are graphed; EC50 values with 95% CI are presented in the tables).

**Table 1.**
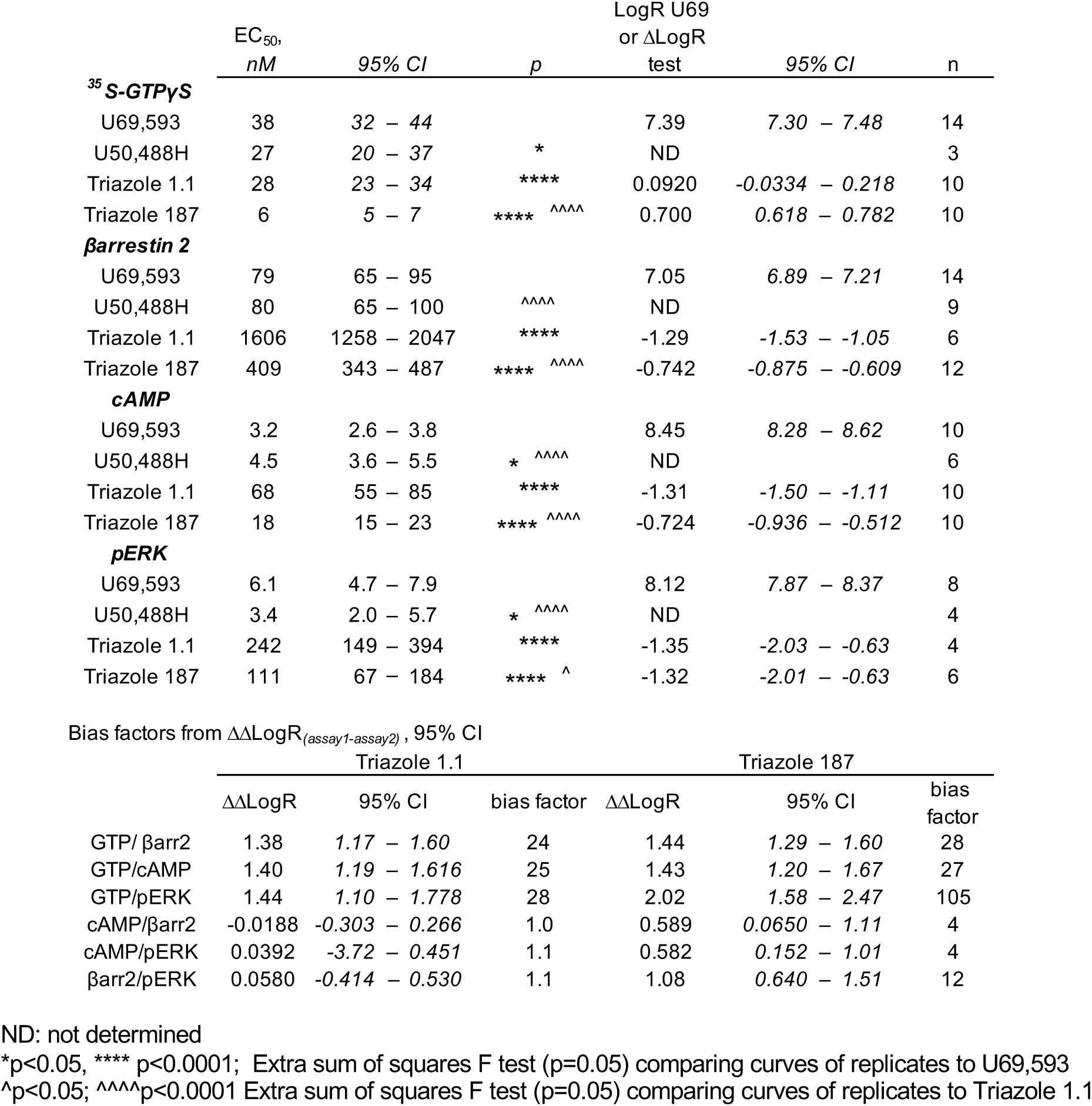
Pharmacological parameters from curve fitting of Figure 1 A-D.

#### Calculation of signaling bias

Bias of the test ligands was determined as previously described^32, 40^. This form of bias analysis employs a reference ligand that is assumed to be a full neutral agonist (i.e., the reference agonist activates all response pathways equally and does not exhibit a preference for one pathway over another). In the experiments presented here, U69,593 was used as the reference agonist; the test agonist and U69,593 curves were fit to the equation^46^:

Reference Agonist

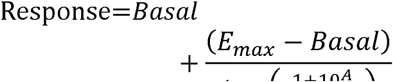

Test Agonist

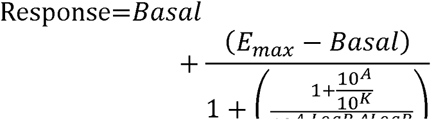

where Basal and Emax are the baseline and maximum response of the system. A is the Log[Agonist], M and K is the equilibrium affinity constant. LogR is the transduction coefficient of the reference agonist (equivalent to the antilog of the reference agonist EC_50_) and ΔLogR is the difference between the LogR of the Reference agonist and the LogR of the test agonist. Using this formulation, it is possible to compare the ΔLogR of a test agonist across different assays to determine if the test agonist exhibits preference for or against a response. In this approach, the ΔLogR values were determined for each experiment and these values were averaged to generate a mean ΔLogR (Stahl et al, 2015). ΔΔLogR values were calculated by subtracting the mean ΔLogR from assay 2 (βarrestin assay) from the similarly calculated mean ΔLogR from assay 1 (GTPγS binding or cAMP inhibition) and is presented as the mean with the 95% confidence intervals. All calculations were performed with GraphPad Prism 10 software; ΔΔLogR values were essentially calculated by an unpaired t test. Bias factors were calculated by taking the antilog of the ΔΔLogR^46, 47^:

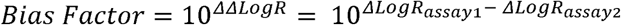

### Statistical and data analyses (*in vivo*)

For most of the behavioral studies, vehicle and U50,448H treatments serves as controls and statistical comparisons are indicated in the figure legends. For determination of potency, data are normalized using the vehicle response and the maximum response to U50,488H to generate the maximum possible effect (%MPE). For the normalization of center time and climbing, the maximum doses of U50,488H were excluded for normalization and nonlinear regression as they were sedating; as indicated in Table 2. Dose response curves of the %MPE data are fit to a hyperbolic function using GraphPad Prism, wherein 100% is used to constrain the maximum fit of the curve. Potency calculations (ED_50_) and the comparison of the highest doses (%MPE) are presented in the table with 95% confidence intervals with statistical analysis. Comparison of potency values are determined by applying an extra sum-of-squares F test (p=0.05) between two curves within the nonlinear regression analysis using Graphpad Prism; comparisons of %MPE in the table are made by ordinary one-way ANVOA. In the figures, raw data are presented as the individual animals and with means ± S.E.M.; effects are compared to vehicle by one-way ordinary ANOVA with a Dunnett’s post-hoc comparison; comparison between drugs is made by Tukey post-hoc comparison, significance was set at p<0.05. Animal numbers are indicated in the figure legends.

**Table 2:** Potency in male mouse behavior assays in Figures 2-4.

|  | **ED<sub>50</sub>,**<br>**mg/kg** | **95% CI** | **p vs**<br>**U50** | **p vs**<br>**1.1** | **% MPE** | **95% CI** | **p vs**<br>**U50** |
| --- | --- | --- | --- | --- | --- | --- | --- |
| **Pruritis** |  |  |  |  |  |  |  |
| U50,488H | 0.17 | 0.086 – 0.36 |  |  | 100 | 84 – 115 |  |
| Triazole 1.1 | 0.58 | 0.30 – 1.11 | ** |  | 73 | 50 – 95 |  |
| Triazole 187 | 0.16 | 0.027 – 0.69 |  | * | 78 | 64 – 93 |  |
| **Diuresis** |  |  |  |  |  |  |  |
| U50,488H | 7.6 | 4.8 – 12 |  |  | 100 | 53 – 146 |  |
| Triazole 1.1 | 87 | 50 – 188 | **** |  | 16 | -3 – 35 | *** |
| Triazole 187 | 57 | 32 – 122 | **** | ** | 23 | 0 – 45 | ** |
| **Climbing<sup>^</sup>** |  |  |  |  |  |  |  |
| U50,488H | 1.3 | 0.33 – 3.5 |  |  | 100 | 42 – 158 |  |
| Triazole 1.1 | 41 | 21 – 91 | **** |  | 107 | 19 – 105 |  |
| Triazole 187 | 17 | 8 – 46 | **** |  | 36 | 6 – 66 |  |
| **Nesting** |  |  |  |  |  |  |  |
| U50,488H | 0.85 | 0.45 – 1.4 |  |  | 100 | 90 – 109 |  |
| Triazole 1.1 | 25 | 13 – 49 | **** |  | 85 | 48 – 122 |  |
| Triazole 187 | 9.2 | 4.7 – 19 | **** | * | 84 | 32 – 137 |  |
| **Sedation (distance)** |  |  |  |  |  |  |  |
| U50,488H | 5.4 | 3.0 – 10 |  |  | 100 | 88 – 112 |  |
| Triazole 1.1 | nc |  |  |  | 10 | -38 – 57 | ** |
| Triazole 187 | 76 | 35 – 267 | **** |  | 29 | -28 – 87 | * |
| **Sedation (beam)** |  |  |  |  |  |  |  |
| U50,488H | 5.2 | 3.3 – 8.4 |  |  | 100 | 93 – 106 |  |
| Triazole 1.1 | nc |  |  |  | 6 | -35 – 46 | *** |
| Triazole 187 | 131 | 64 – 489 | **** |  | 19 | -31 – 70 | ** |
| **Center Time<sup>^</sup>** |  |  |  |  |  |  |  |
| U50,488H | 1.01 | 0.11 – 3.6 |  |  | 100 | 46 – 154 |  |
| Triazole 1.1 | nc |  |  |  | 5 | -35 – 45 | ** |
| Triazole 187 | 3.6 | 1.5 – 7.6 |  |  | 65 | 25 – 105 |  |
| **Center Distance** |  |  |  |  |  |  |  |
| U50,488H | 5.4 | 2.9 – 10 |  |  | 100 | 88 – 111 |  |
| Triazole 1.1 | nc |  |  |  | 12 | -45 – 70 | ** |
| Triazole 187 | 53 | 25 – 146 | **** |  | 37 | -16 – 90 | * |

\*p<0.05, \*\*p<0.01; \*\*\*p<0.001 \*\*\*\* p<0.0001; Extra sum of squares F test (p=0.05) comparing the normalized %MPE curves.
<sup>^</sup> for Climbing, the highest point used is 5 mg/kg U50,488H; for Center time, the highest point used is 15 mg/kg U50,488H for normalization.

## RESULTS

### Pharmacological and pharmacokinetic characterization of Triazole 187

Triazole 187 shows improved affinity over triazole 1.1 and U69,593, in [^3^H]-Naloxone competition binding assays (pK*_i_*± S.E.M., M: U69: 9.15 ± 0.084; Tri1.1: 9.03 ± 0.084; Tri187: 9.63 ± 0.13; compared to triazole 187: *p*=0.0002 vs Tri1.1; *p*=0.0027 vs U69, n=3) (SFig. 1A). Triazole 187 is more potent than triazole 1.1 and U69,593 for promoting [^35^S]-GTPyS binding, inhibiting forskolin-stimulated cAMP accumulation, and recruiting βarrestin2 to KOR (Fig. 1B-D, Table 1). In ERK1/2 phosphorylation, triazole 187 does not gain potency relative to triazole 1.1 but retains poor potency relative to U69,593 (Fig. 1E, Table 1). To compare the relative potency between the assays, a bias factor analysis was undertaken, using U69,593 as the reference agonist, as previously described^22, 32, 48^ (Table 1, Fig. 1F). While the potency of triazole 187 improves, the overall bias for GTPγS binding over the other signaling assays remains (Table 1).

**Figure 1.**
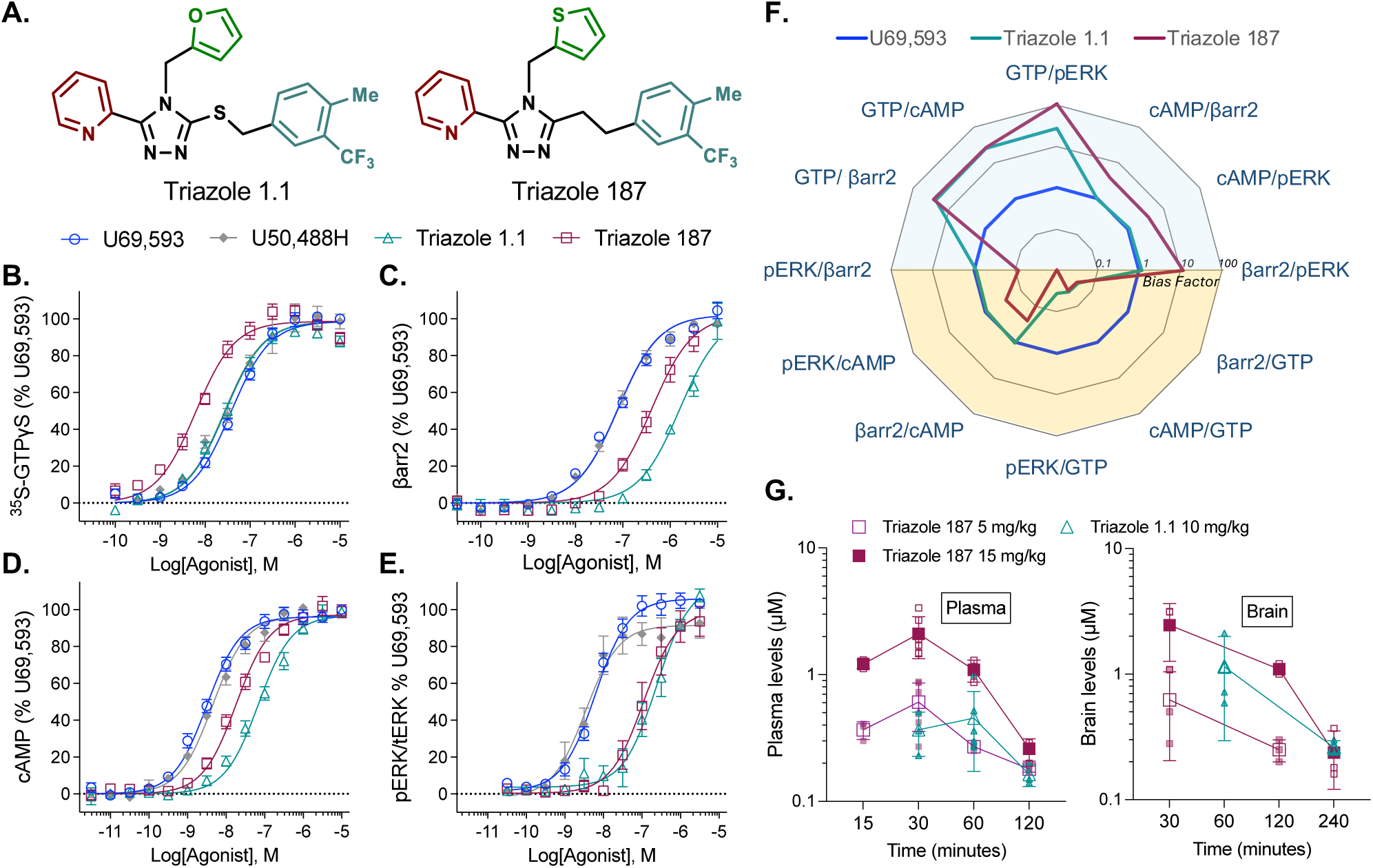
Pharmacological and pharmacokinetic characterization of triazole 187 compared to U50,488 and triazole 1.1. **(A)** Structures of triazole 1.1 and triazole 187. **(B)** ^35^S-GTPyS binding; **(C)** βarrestin2 recruitment; **(D)** inhibition of forskolin-stimulated cAMP accumulation; and **(E)** ERK1/2 phosphorylation are presented as % maximal U69,593 response. Data are presented as mean ± SEM; potencies with 95% CI and *n* are detailed in Table 1. **(F)** Bias factors were calculated using U69,593 as the reference agonist, which defines unity. Bias factors are plotted on a logarithmic scale (base 10) (10^ΔΔLogR) in the web plot; as presented, a value of >10 indicates a preference for the numerator. **(G)** Pharmacokinetic properties of triazole 187 compared to triazole 1.1 measured in C57BL6/J male mice: shown are plasma (left) and brain (right) levels after intraperitoneal injection (i.p.) over time at the doses indicated presented as mean ± SD (*n* = 3 brains; *n* = 3 plasma except *n* = 6 triazole 1.1 at 60 min and *n* = 6 for triazole 187 at 30 min).

Triazole 187 was tested for off-target binding at 42 potential receptor targets by the NIMH-Psychoactive Drugs Screening Program (PDSP) which, did not reveal any high affinity targets other than KOR (SFig. 1B). Triazole 187 was also evaluated for brain and plasma levels in C57BL6/J mice (Fig 1G); like triazole 1.1 (which was first published in ^22, 30^ and repeated here); triazole 187 is also brain penetrant. Triazole 187 can still be detected in brain at 4 hours.

### Triazole 187, triazole 1.1, and U50,488H are potent anti-pruritic compounds

Triazole 1.1 was previously shown to be equally efficacious as U50,488H in the chloroquine phosphate-induced pruritis mouse model when compared at a dose of 1 mg/kg^22^. Here, we show that triazole 187 is equiefficacious and equipotent to U50,488H while triazole 1.1 is less potent than both compounds (Fig. 2A, Table 2). U50,488H and triazole 187 suppresses scratching at 0.3 mg/kg, while only the highest (1.0 mg/kg) dose of triazole 1.1 produces a significant effect (Fig. 2B).

**Figure 2.**
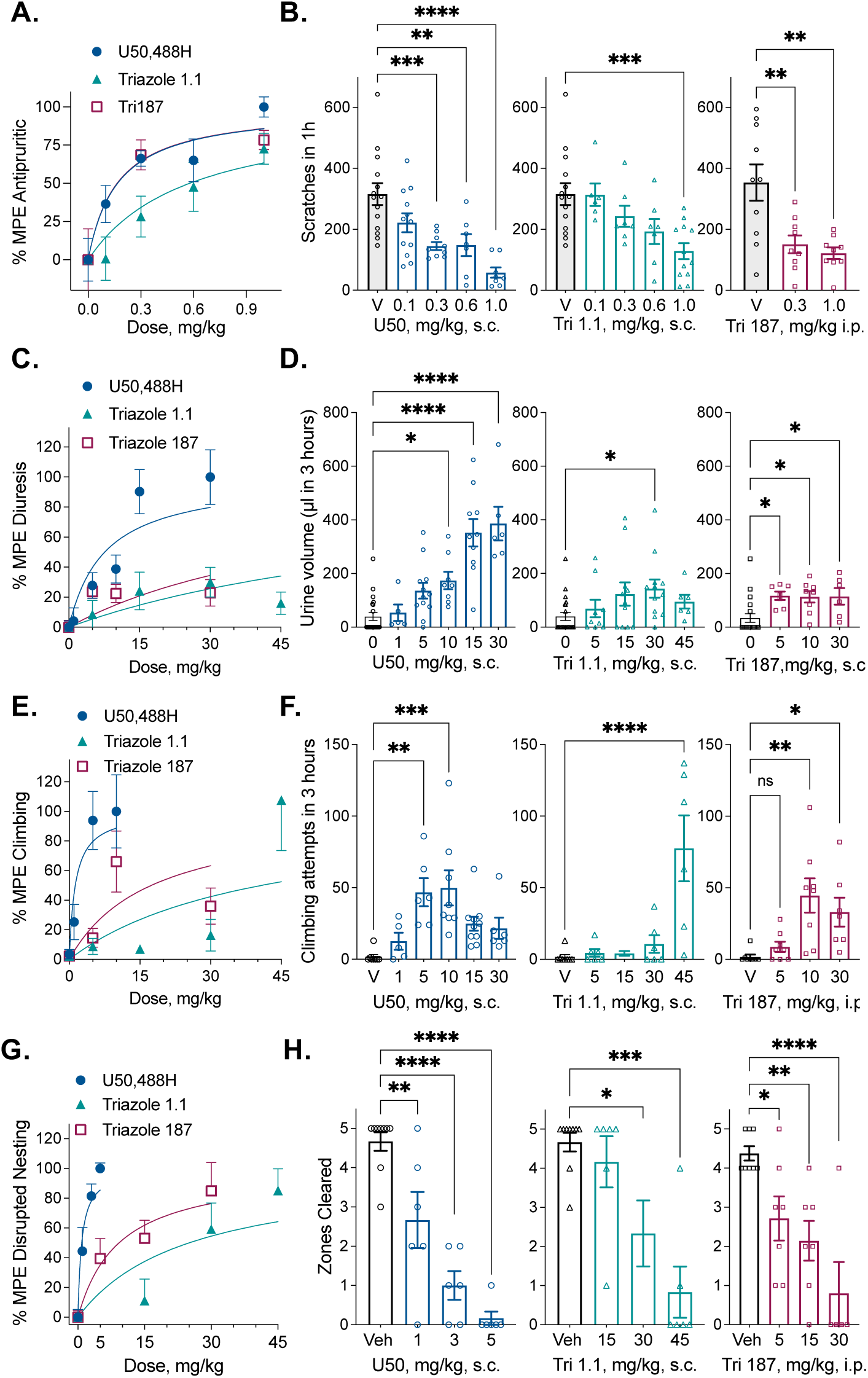
KOR agonists effects on chloroquine phosphate-induced pruritis diuresis, climbing behavior and nesting behavior in male C57BL6/J mice. **(A)** U50,488H, triazole 1.1, and triazole 187 dose-dependently suppress chloroquine-phosphate (CP) (40 mg/kg, s.c. neck) induced scratching in in male C57BLJ/6 mice when administered 10 min before CP. **(B)** The number of scratches are shown (*n*: Veh s.c.: 14; U50: 7-12; Tri 1.1: 6-12; Veh i.p.: 10 Tri 187: 9). **(C)** U50,488H, induces more urine production than triazole 1.1 and triazole 187 over a 3 hour period comparing. **(D)** The urinary output in µL is shown per dose (*n*: Veh s.c.: 21; U50: 5-12; Tri 1.1: 6-12; Tri 187: 6-8). **(E)** Climbing responses were counted during the urine collection over 3 hours. Since U50,488H is sedative at higher doses, the 10 mg/kg dose was used to set the maximum and 15 and 30 mg/kg data were not fit to determine potency in the %MPE curve. **(F)** Climbing attempts are shown for each dose (*n* are: the same as in **D**). **(G)** U50,488H potently disrupts nesting behaviors. **(H)** The number of zones cleared are shown for each dose (*n*: Veh s.c.:9; Veh i.p.: 8; U50: 6; Tri 1.1: 6-7; Tri 187: 5-7). The potency (ED_50_) from the hyperbolic curves for each compound is presented in Table 2 with 95% CI. Data are presented as mean ± S.E.M. Dose vs. vehicle comparisons were conducted using ordinary one-way ANOVA with Dunnett’s post-hoc test (\**p<0.05, **p<0.01, ***p<0.001, ****p<0.0001*).

### Triazole 1.1 and triazole 187 produce less diuresis in mice

Diuresis is a common side effect of typical KOR agonists^24^. In mice, U50,488H, triazole 1.1 and triazole 187 increase urine output over a 3-hour period (Fig. 2C, D); however, triazole 1.1 and triazole 187 are significantly less potent and efficacious than U50,488H (Fig. 2D, Table 2). Notably, the total urine volume produced by the highest doses of triazole 1.1 (95 ± 26 µL) and triazole 187 (115 ± 31 µL) is significantly less compared to U50,488H (386 ± 63 µL) (one-way ordinary ANOVA p<0.01, n = 6 per group; Fig. 2D, Table 2) suggesting that while a diuretic effect remains, the overall impact is less pronounced for the two biased agonists.

### U50,488H and triazole 187 induce climbing behavior in mice

While monitoring mice for diuresis, we observed mice attempting to climb out of the assessment box (12 cm wall height); these attempts were recorded over the 3-hour test period. U50,488H was more potent than either triazole for increasing climbing behaviors (Fig. 2E, F, Table 2); higher doses of U50,488H are known to be sedative^22, 23, 49^ therefore, the effects at 15 and 30 mg/kg were not included in determining the potency of U50,488H (Fig. 2E, Table 2).

### Triazole 1.1 and triazole 187 produce less disruption of nesting behaviors than U50,488H

Nest building is an instinctive behavior in mice that can be disrupted by drugs that induce both pleasant and aversive effects in humans, including cannabis, alcohol, morphine and antidepressants^44, 50–54^. KOR agonists have also been shown to disrupt nesting behavior^44, 55^. As anticipated, U50,488H potently disrupts mouse nesting behavior at doses as low as 1 mg/kg (Fig. 2G, H, Table 2). While the triazoles are able to interfere with nesting, they are significantly less potent than U50,488H (Fig. 2G, Table 2).

### Triazole 1.1 and triazole 187 do not induce sedation in mice

In mice, sedative drugs decrease spontaneous locomotor activity in an open-field^52, 56^. Consistent with our prior studies, U50,488H rapidly decreases the total distance traveled in one hour in a dose responsive manner (Fig. 3A, B Table 2). This is recapitulated in measures of horizontal activity as detected by the number of beam breaks in the field (Fig. 3C). Triazole 1.1 and triazole 187 (Fig. 3A-C) do not significantly decrease the distance traveled or horizontal activity at any dose tested relative to vehicle. In female mice, only the highest dose of U50,488H (15 mg/kg) decreases total distance travelled (Fig. 3D), although decreases in the horizontal activity (Fig. 3E) mirrors the effects of U50,488H observed in the males (Fig. 3C). Similar to the males, total distance and horizontal activity in female mice are not decreased by triazole 1.1 or triazole 187 (Fig. 3D, E). The sedating effects of U50,488H are selective for KOR as KOR-KO mice do display this effect ^22^ (SFig. 2A).

**Figure 3.**
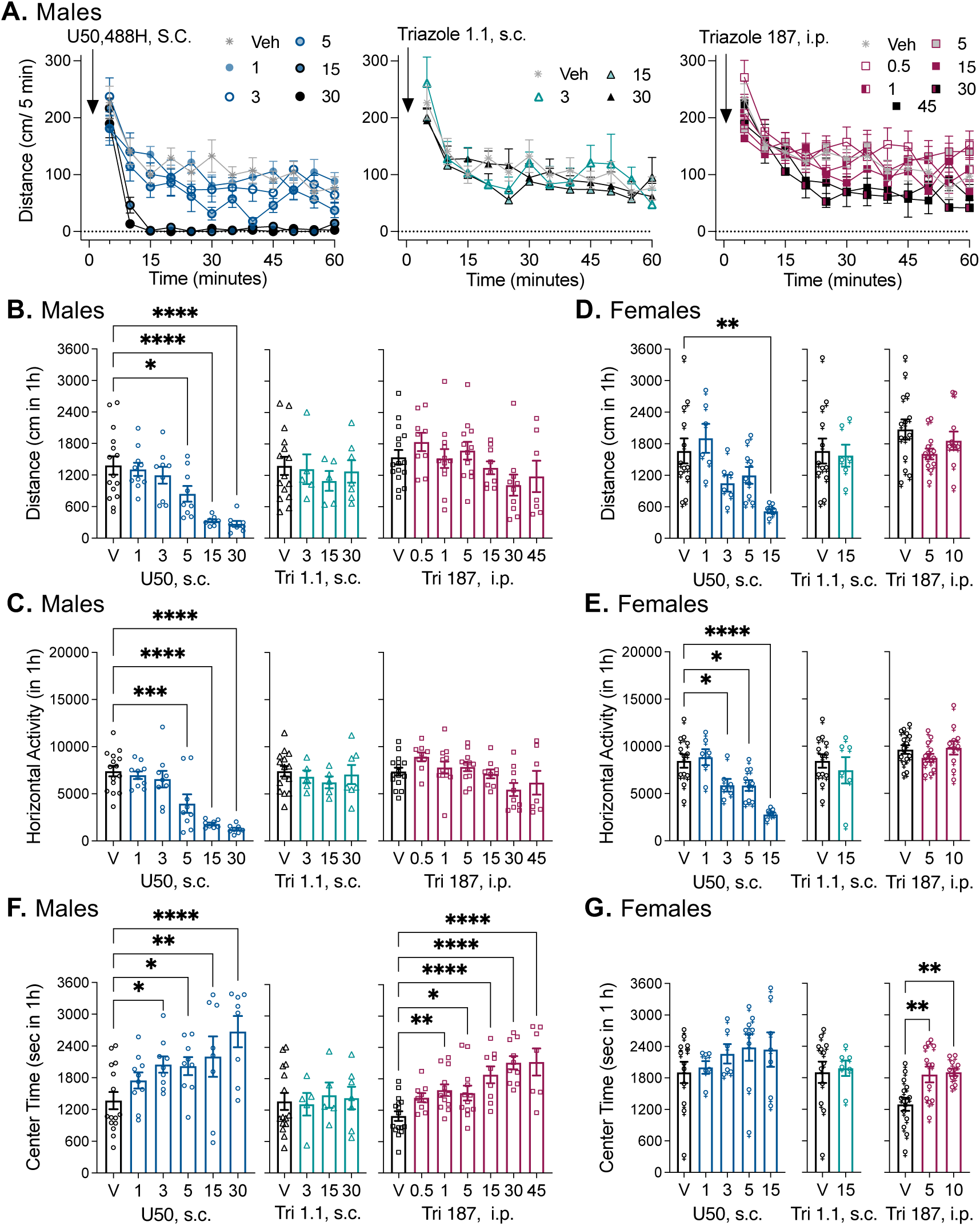
Triazole 187 increases center time but does not induce sedation in male and female C57BL6/J mice. **(A)** The distance traveled in male C57BL6/J is rapidly decreased by U50,488H while triazole 1.1 and triazole 187 do not produce this effect over time (time x dose, 2-way RM-ANOVA U50,488H: F_(55, 583)_ = 1.797, *p*=0.0006; triazole 1.1: F_(33, 308)_ = 0.7777, *p*=0.8068; triazole 187: F_(66, 748)_ = 1.279, *p*=0.0734). **(B)** The sum of the total distance traveled over 1 hour and **(C)** the sum of horizontal activity (beam breaks) were significantly decreased by U50,488H treatment, while triazole 1.1 and triazole 187 have no effect compared to vehicle in male C57BL6/J mice. **(D)** In female mice, a similar effect was observed for total distance and **(E)** horizontal beam breaks. **(F)** In male mice, U50,488H increases total time spent in the center of the open field over 1 hour compared to vehicle; triazole 1.1 has no significant effect, while triazole 187 also dose-dependently increases center time. **(G)** U50,488H and triazole 1.1 do not significantly increase center time in the female mice; however, triazole 187 increases center time. Data are presented as mean ± S.E.M. and potencies are presented with 95% CI in Table 2. (males: *n* = Veh s.c., 15; Veh i.p., 16; U50, 8-10; Tri 1.1, 5-7; and Tri 187, 7-12; females: *n* = Veh s.c.,12; Veh i.p., 14; U50, 6-9; Tri 1.1, 6; and Tri 187, 10-12). Dose vs. vehicle comparisons were conducted using ordinary one-way ANOVA with Dunnett’s post-hoc test (\**p<0.05, **p<0.01, ***p<0.001, ****p<0.0001*). Triazole 1.1 and U50,488H were administered s.c. and their vehicle is given via the same route. Triazole 187 and its vehicle were administered by the i.p. route. Potency (ED_50_) values with 95% CI are presented in Table 2. The potency of U50,488H in female mice could be calculated only for total distance and beam breaks and are 4.9 *(2.4-10)* and 4.5 *(2.7-7.6)* mg/kg, s.c., respectively.

### Triazole 187 and U50,488H increase time spent in the center of the open field

The open field activity measures were also analyzed for time spent in the center of the field, as an indicator of anxiolytic-like behavior in mice, based on the animaĺs innate aversion to novel open areas^57^. In males, U50,488H increases center time in a dose-dependent manner (Fig. 3F, Table 2); although movement within the center zone is significantly decreased at 5, 15 and 30 mg/kg (SFig. 2B, Table 2), mirroring the total distance traveled (Fig. 3A, B). Triazole 1.1 does not affect the time spent in the center compared to vehicle at any dose tested (Fig. 3F) nor does it affect the distance traveled within the center zone (SFig 2B). In contrast, triazole 187 increases center time in a dose dependent manner (Fig. 3F) without decreasing overall distanced traversed in the center (SFig. 2B, Table 2). The effects of triazole 187 and U50,488H on center time and center distance are absent in KOR-KO mice (SFig. 2A). In female mice, U50,488H-induced increases in center time did not reach statistical significance; however center distance was decreased by U50,488H at 15 and 30 mg/kg (Fig. 3G, SFig. 2C). Recapitulating the results in the males; triazole 1.1 produced no changes in center time or center distance in female mice and triazole 187 significantly increased center time in females without decreasing center distance (Fig. 3G, SFig. 2C).

### Triazole 187 increases time spent in open arms of the elevated plus maze

An elevated plus maze was used to further investigate anxiolytic-like behaviors in male (Fig. 4A) and female (Fig. 4B) mice. In males, when tested at a low dose to avoid sedation, U50,488H does not significantly increase time spent in the open arms. Although U50,488H decreases the total number of arm entries, it does not decrease total distance traversed. Triazole 1.1, tested at 15 mg/kg, has no effect on any other parameter measured; however, Triazole 187 (5 mg/kg) increases the time spent in the open arms and in the distal end of the open arms without affecting arm entries or overall activity (Fig 4A). In female mice, 5 mg/kg of triazole 187 does not significantly increase time spent in the open arms or in the distal end of the open arms. However, female mice enter the arms more frequently and cover greater distance in the 5-minute test with triazole 187 treatment compared to vehicle (Fig. 4B).

**Figure 4.**
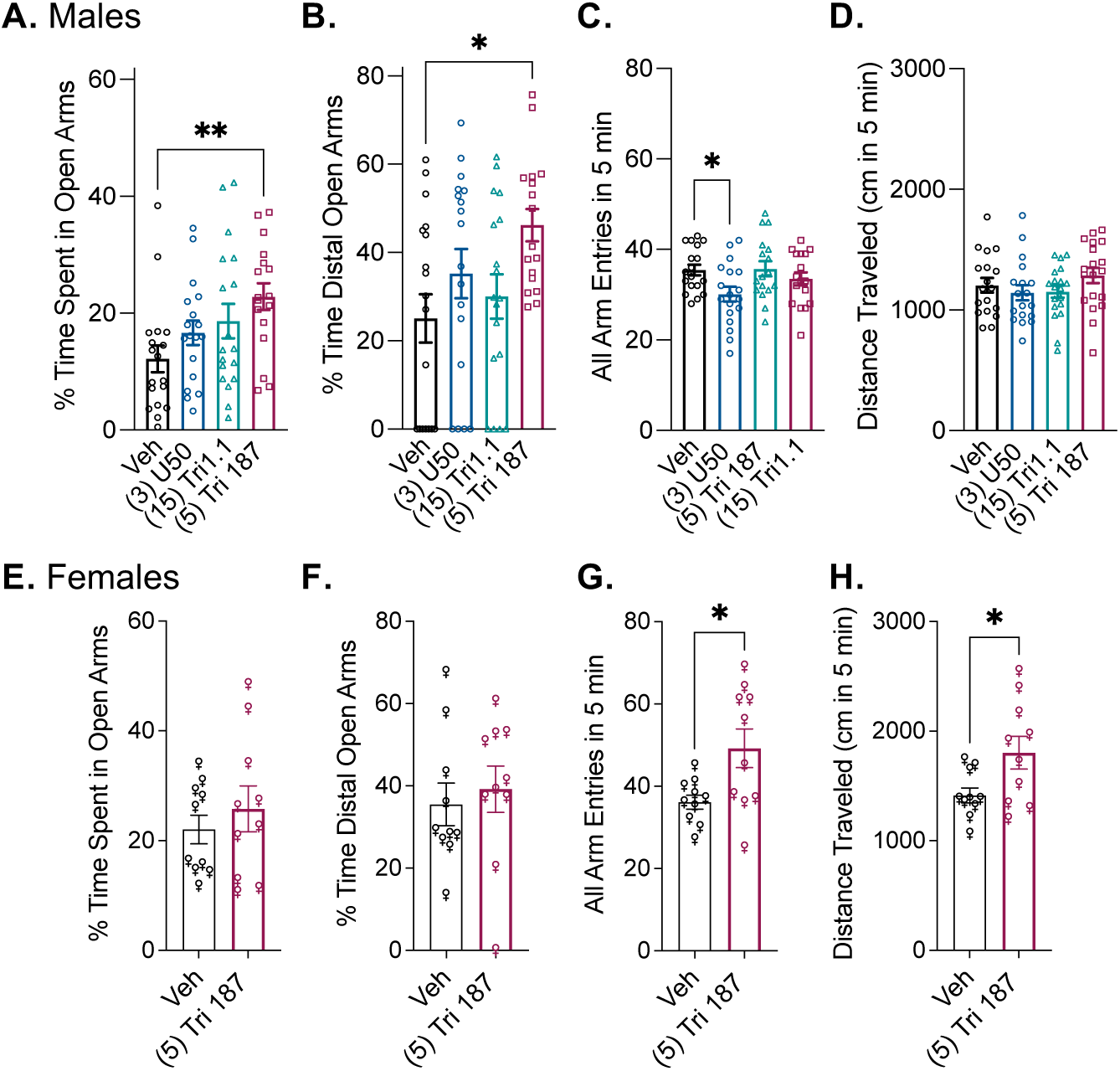
Triazole 187 increases time spent in open arms of the elevated plus maze in male mice and increases all arm entries and distance travelled in female mice. Vehicle (i.p.), U50,488H (3 mg/kg, i.p.), triazole 1.1 (15 mg/kg, i.p.) or triazole 187 (5 mg/kg, i.p.) were administered to male mice **(A)** and vehicle or triazole 187 (5 mg/kg, i.p.) were given to **(B)** female mice 30 minutes prior to the start of the 5-minute test which measures the percent of time spent in the open arms; the percent of time spent in the distal part of the open arm; the total number of arm entries; the total distance travelled in the entire maze in 5 minutes. Data are presented as mean ± S.E.M. (A) Males: *n* = Veh i.p.: 18; U50: 18; Tri 1.1: 18; and Tri 187: 17; drug vs vehicle: (*p<0.05, **p<0.01 by one-way ordinary ANOVA, with Dunnett’s post-hoc test). (B) females: *n* = Veh i.p.: 10; and Tri 187: 10; triazole 187 compared to vehicle: (*p<0.05 unpaired t-test).

## DISCUSSION

Triazole 187 is a member of a new series of compounds recently reported by our group^37^ that shows improved affinity, potency and preserved selectivity and bias for promoting GTPγS binding to KOR over several other signaling pathways. In mice, triazole 187 is more potent in suppressing chloroquine phosphate-induced itch than triazole 1.1, while retaining a lack of sedation, suggesting that potency can be improved while preserving the widened therapeutic window. While diuresis is induced by all the agonists tested, the overall volume produced by triazole 1.1 and triazole 187 is less than half as much as U50,488H, suggesting that this adverse effect may also be diminished for these agonists. Unlike U50,488H, triazole 187 increases time spent in the center region of the open field while not producing sedation, potentially indicating anxiolytic-like effects of the biased KOR agonist.

Clinically, KOR agonists are prescribed for the treatment of non-histamine pruritis that can accompany hemodialysis^3^. However non-dermatitis, non-inflammatory itch is difficult to treat; moreover, its condition has been described as tormenting to endure, leading to severe impacts on mood. Chronic itch often produces anticipation and anxiety in patients about future symptoms, which can contribute to the worsening of the condition^58, 59^. Extensive studies show that psychiatric comorbidities, including anxiety and depression, are commonly found in patients with chronic pruritus. These patients also experience high rates of sleep disturbances and report a significantly lower quality of life^60–64^.Our results present a promising strategy, by CNS-permeable biased KOR agonists, for advancing the treatment of pruritus and the psychological comorbidities that can accompany this condition.

It is known that activation of KOR in the CNS produces psychoactive effects^10,65,,66^ and modulates various mood states, such as anxiety and depression^10, 35, 67–70^. KOR antagonists have been pursued for treating neuropsychiatric conditions, with recent advancements in drug development aimed at addressing stress-related mood disorders and major depressive disorders^71–73^. In contrast, conventional KOR agonists are generally not considered viable clinical interventions due to the adverse effects associated with their use. However, a biased KOR agonist may present a new means to modulate mood without sedation and may provide a means to treat pain and itch with while alleviating anxiety associated with ongoing itch.

Prior studies have shown that the conventional KOR agonists, such as U50,488H, produce anxiolytic-like effects in rodents in the elevated plus maze^12^, but since assessments of anxiolytic-like behaviors in mice measure movement, the sedative effects induced by these compounds can confound interpretation^74, 75^. In this study, upon using lower doses of U50,488H (1 and 3 mg/kg) to avoid sedation, we were able to observe increases in center time, but not increases in time spent in the open arms of the elevated plus maze. Conversely, triazole 187, which does not induce sedation, increases in center time and demonstrates efficacy in the elevated plus maze, potentially indicating anxiolytic-like effects. Notably, these drug effects can be observed in the same cohort of mice after drug treatment, as both the open-field activity and elevated plus maze simultaneously measure the overall distance traveled and time spent in the center of the open-field or the open arms of the maze. Taken together, we demonstrate that the diverging behavioral effects of a biased agonist can be differentiated within the same cohort of animals in a dose dependent manner; we show that anxiolytic behaviors can be enhanced without sedation.

An increase in climbing behavior was also observed in mice induced by sub-sedative doses of U50,488H, as well as triazole 187. Enhanced climbing behavior has previously observed with U69,593 in mice^76^ and could indicate novelty seeking and exploring^77^. On the other hand, climbing behavior may be representative of drug-induced escape behavior^78^. In this study, we also found that U50,488H disrupts mouse nesting behavior at sub-sedative doses (1 and 3 mg/kg), while triazole 187 and triazole 1.1 are significantly less potent than U50,488H (Table 2). Nesting behavior can be disrupted by both positive and negative stimuli^44, 50–54^, making it challenging to fully interpret these results as favorable or aversive^79^. Regardless, there remains a significant reduction in potency for the biased agonists relative to U50,488H in the nesting and the climbing studies (Table 2).

Previous studies have shown that female rodents can be less responsive to KOR agonists in behavioral models of mood disturbances. For example, decreased responses to the dysphoria-like effects of U50,488H, were noted in female rats in intracranial self-stimulation (ICSS) studies^80, 81^ and female mice were less responsive to U50,488H in anxiety-related behaviors and conditioned place aversion (CPA) in female mice^55, 82^. We find that while U50,488H dose-dependently decreases horizontal activity in the open-field in female mice, we did not observe a significant increase in center time or in the choice of the open arms in the elevated plus maze. However, triazole 187 retains efficacy in both male and female mice for increasing the time spent in the center area while it produces no sedation in either sex. In the elevated plus maze, triazole 187 does not increase time spent in the open arms in female mice; however, they do show increases in the total distance traversed the total arm entries which could be the manifestation of exploratory behaviors. Taken together, triazole 187 shows favorable anxiolytic-like effects in both male and female mice without the added confound of sedation.

Aside from sedation, another disruptive side effect of KOR agonists is diuresis, which occurs in both mice and humans^24, 83–85^ Moreover, the clinically used difilikefalin, despite its peripheral restriction, still induced significant diuresis which is mediated by actions in the CNS^86^. Additionally, nalfurafine has been reported to induce diuresis in rats^87^, while U50,488H has been reported to induce diuresis in rats, as well as mice and non-human primates^24, 49, 84, 88^. Triazole 1.1 and 187 induce a significant increase in diuresis, however, the output is significantly less (∼30%) than that induced by U50,488H. This mild diuretic effect appears to approach a ceiling effect and could be seen as clinical advantage in developing a mild diuretic or as analgesics for pain accompanied by edema.

Overall, this study highlights and contributes to the growing literature demonstrating that it is possible to separate therapeutic benefits from side effects by imparting selectivity for different receptor active states. Triazole 187 represents a new generation of G protein coupling-biased KOR agonists that may improve the future development of non-sedating antipruritics agents and anxiolytics. Additionally, CNS-permeable biased KOR agonists, such as triazole 187, present a promising alternative therapeutic approach for advancing the treatment of chronic itch as well as the anxiety often associated with this condition.

## Declarations

The authors have no competing interests to declare. UF and UNC-CH have filed for patent protection for Triazole 187 and hold a patent on triazole 1.1.

