## Supplemental for "Triazole 187 is a biased KOR agonist that suppresses itch without sedation and induces anxiolytic-like behaviors in mice"

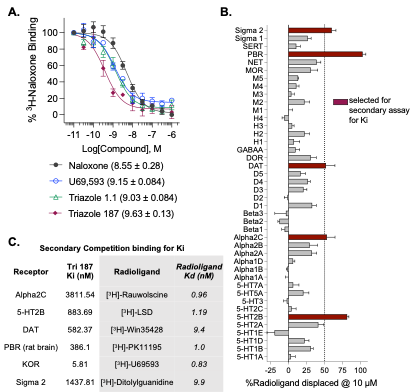

**Figure S1. KOR affinity and selectivity in psychoactive drug target screening of Triazole 187**. **(A)** Binding affinity comparing U69,593, Triazole 1.1, Triazole 187 and naloxone by competition binding using 2 nM ^3^H-Naloxone as the radioligand (the pK*_D_* for naloxone and the pK*_i_* with S.E.M. are presented in the figure). **(B)** The Psychoactive Drugs Screening Program (PDSP) tested the ability of Triazole 187 to displace radioligand binding of 42 psychoactive drug target proteins at a 10 µM concentration. Five receptor targets showed displacement greater than 50% and were further investigated for competition binding assays to determine affinity. **(C)** Binding affinity determined by competition binding. The K*_D_* of the radioligand used in the competition assays is provided for comparison (from the PDSP).

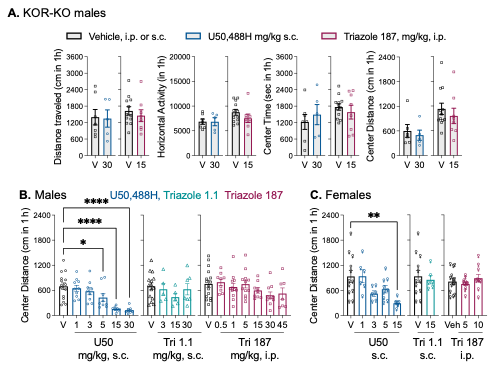

**Figure S2. KOR agonists effects on spontaneous locomotor activity in KOR-KO mice and U50,488H decreases distance travelled in the center in male and female C57BL6/J mice.** **(A)** Distance travelled, horizontal activity, center time and center distance did not differ between vehicle and drug treated mice in KOR-KO mice. **(B)** U50,488H decreased total distance travelled in the center of the open-field over 1-hour compared to vehicle**;** triazole 1.1 and triazole 187 had not effect on center distance travelled.  **(C)** In female C57BL6/J mice U50,488H decreased total distance travelled in the center of the open-field over 1-hour at 15 mg/kg U50,488H compared to vehicle**,** triazole 1.1 and triazole 187 had not effect on center distance travelled. Data are presented as mean ± SEM and potencies are presented with 95% CI in Table 2. (KOR-KO: male *n* = Veh s.c., 7; Veh i.p., 12; U50, 5; Tri 1.1, 7; male C57BL6/J: *n* = Veh s.c., 15; Veh i.p., 16; U50, 8-10; Tri 1.1, 5-7; and Tri 187, 7-12; female C57BL6/J: *n* = Veh s.c.,12; Veh i.p., 14; U50, 6-9; Tri 1.1, 6; and Tri 187, 10-12). Drug vs. vehicle comparisons were conducted using ordinary one-way ANOVA with Dunnett’s post-hoc test (**p<0.05, **p<0.01, ***p<0.001, ****p<0.0001*). Triazole 1.1 and U50,488H were administered s.c. and their vehicle is given via the same route. Triazole 187 and its vehicle were administered by the i.p. route. Potency (ED_50_) values with 95% CI for center distance are presented in Table 2 for male mice. For female mice, the potency of U50,488H is 3.5 (1.7 –7.2) mg/kg, i.p..
